# Immediate-Early Genes as Influencers in Genetic Networks and their Role in Alzheimer’s Disease

**DOI:** 10.1101/2024.03.29.586739

**Authors:** Margarita Zachariou, Eleni M. Loizidou, George M. Spyrou

**Affiliations:** Bioinformatics Department, The Cyprus Institute of Neurology and Genetics; biobank.cy, Center of Excellence in Biobanking and Biomedical Research, University of Cyprus

**Keywords:** Immediate Early Genes, Activity Regulated Genes, Genetic Networks, Alzheimer’s Disease, *MARK4*, GWAS, eQTL

## Abstract

Immediate-early genes (IEGs) are a class of activity-regulated genes (ARGs) that are transiently and rapidly activated in the absence of de novo protein synthesis in response to neuronal activity. We explored the role of IEGs in genetic networks to pinpoint potential drug targets for Alzheimer’s disease (AD). Using a combination of network analysis and genome-wide association study (GWAS) summary statistics we show that (1) IEGs exert greater topological influence across different human and mouse gene networks compared to other ARGs, (2) ARGs are sparsely involved in diseases and significantly more mutational constrained compared to non-ARGs, (3) Many AD-linked variants are in ARGs gene regions, mainly in *MARK4* near FOSB, with an AD risk eQTL that increases *MARK4* expression in cortical areas, (4) *MARK4* holds an influential place in a dense AD multi-omic network and a high AD druggability score. Our work on IEGs’ influential network role is a valuable contribution to guiding interventions for diseases marked by dysregulation of their downstream targets and highlights *MARK4* as a promising underexplored AD-target.

**Highlights:**

- Immediate-early genes are topologically influential in brain genetic networks in mouse and human.
- Activity-regulated Genes (ARGs) are highly constrained with sparse gene-disease relevance.
- There are several AD-associated variants in ARGs gene regions, mainly in *MARK4* near *FOSB*.
- GWAS and network analysis of ARG’s pinpoint *MARK4* as a promising underexplored AD target.

## 1. Introduction

Neuronal activity elevation induces the expression of several activity-regulated genes (ARGs)^1^. Activity-dependent regulation of gene expression has emerged as a prevailing explanatory mechanism of how molecular-level activity translates to neuron-level functions pertinent to neuronal connectivity and synaptic plasticity^2^. ARGs can be divided into two groups based on their response time, namely primary response genes (PRGs) and secondary response genes (SRGs)^3^. PRGs are rapidly induced without the need for de novo translation/protein synthesis whereas SRGs induction has a slower onset, requires de novo protein synthesis and is regulated by PRGs^3,4^. PRGs can be further classified into rapid primary response genes (rPRGs) and delayed primary response genes (dPRGs) according to their peak response time^5^. Different patterns of neuronal activity can evoke a different gene expression program; brief neuronal activity selectively evokes the response of the first of the three temporal waves (Figure 1A) and requires MAPK/ERK signalling while sustained neuronal activity is required to elicit the response of all three transcriptional waves of rPRGs, dPRGs and SRGs^3,5^. Rapid PRGs that are transiently and rapidly activated in cells in response to a variety of cellular stimuli are also known as immediate-early genes (IEGs). In neurons, IEGs are involved in regulating key processes such as synapse formation, transmission, and plasticity^6^. Deregulation of neuronal IEGs is associated with the onset of various brain disorders, including neuropsychiatric and neurodegenerative diseases^6,7^. IEGs, whose expression levels reflect recent changes in neuronal activity, are often used as proxies for when and where neuronal activity took place, especially in difficult-to-access regions, with live imaging ^1,8–10^. Beyond their key role in neuronal circuits, IEGs are also involved in critical roles in other tissues^11^ and are thus considered less amenable as pharmacological targets for the treatment of brain diseases. A possible mitigation has been proposed to uncover upstream modulators and downstream targets of IEGs that could be more amenable to manipulation by pharmacological or other means ^2^. To that end, an extended outstanding question is to pinpoint the targets/modulators of IEGs in specific neuronal cell types and specific brain circuits^12^.

**Figure 1:**
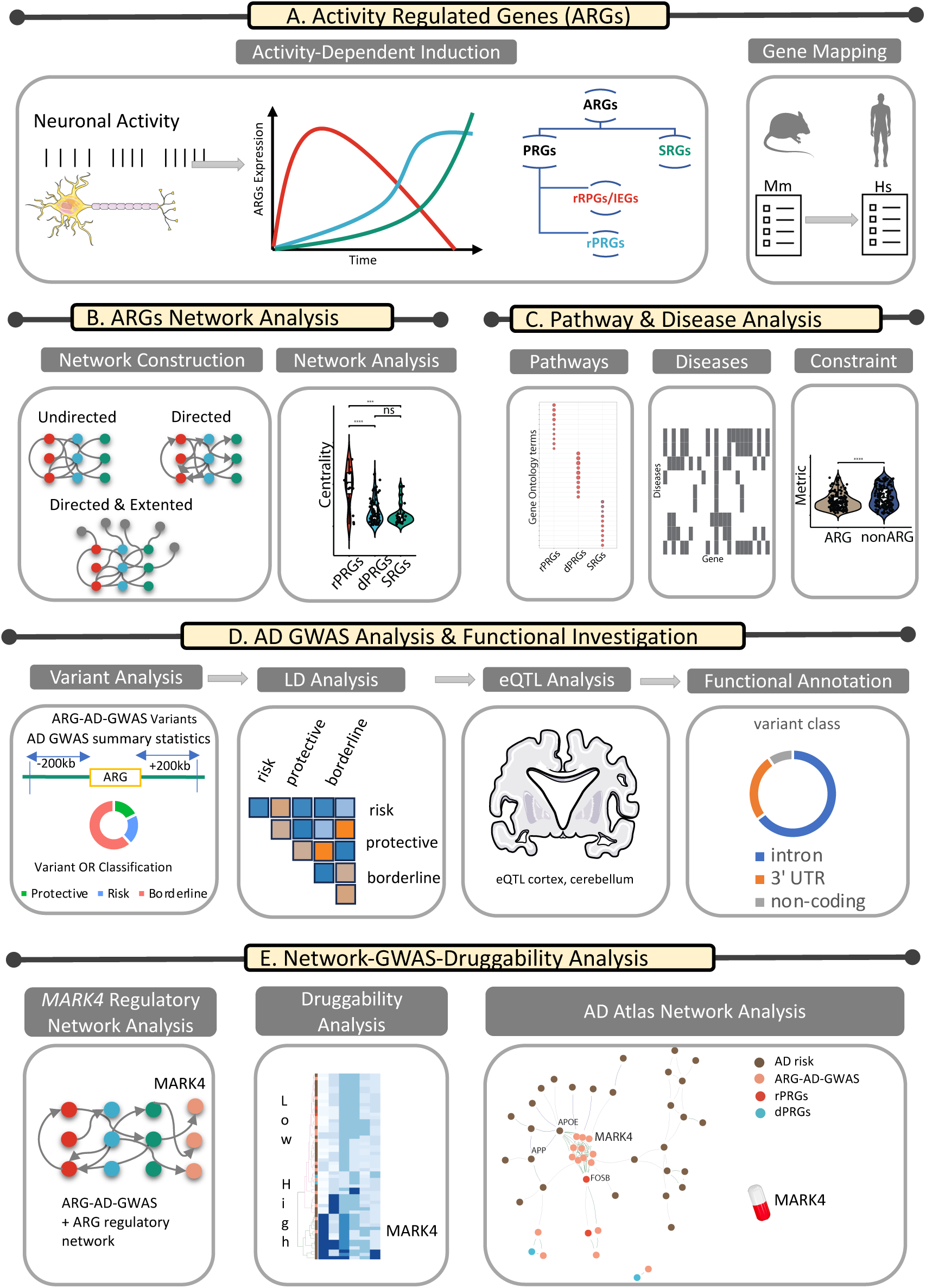
Illustration of the workflow. More details are available in the text.

Motivated by the knowledge gap on how IEGs interact at the network level and whether such a systemic approach can benefit our investigation of drug targets for brain diseases, we set out to explore the interaction and topological influence of IEG as part of genetic ARG network ensembles and their relevance to Alzheimer’s Disease (AD).

AD is the most prevalent neurodegenerative disease posing a tremendous economic and human burden on society worldwide^13^. Pathological hallmarks of AD include the presence of amyloid plaques and neurofibrillary tangles in the brain. Aggregation of amyloid beta (Aβ) initiates a cascade of events that lead to the formation of amyloid plaques and neurofibrillary tangles, which ultimately cause neuronal injury or loss in particular areas of the brain^14^. IEGs are involved in key neuronal functions, such as synaptic plasticity and memory, known to be impaired in the brain in AD and other brain diseases. Research indicates that one of the key players in AD pathology, the amyloid precursor protein (*APP*), regulates epigenetically the expression of PRGs (*Egr1*, *c-Fos*, *Bdnf*, and *Arc*) known to be involved in memory formation^15^. IEGs are potentially also relevant to AD pathology due to their activation in response to environmental stimuli, including stress ^16,17^ – a known risk factor for AD ^18,19^.

This paper is organized as follows, as illustrated in the workflow in Figure 1: (1) We use a mouse ARG data set and its mapped human genes to investigate whether IEGs also share key topological properties beyond their gene-induction mode that separate them from other ARGs (Figure 1A and 1B). (2) We investigate the role of ARGs in biological pathways, diseases and in terms of their mutational constraint profile (Figure 1C). (3) With AD as a case study, we explore in detail the role of variants in or close to ARGs with genome-wide association study (GWAS) summary statistics and functional investigation (Figure 1D), (4) by combining the network approach and the GWAS annotated ARGs in AD, we highlight genes close to ARGs as potential drug targets for AD, with a focus on *MARK4*, using a combination of network theory approach and GWAS summary statistics analysis (Figure 1E).

## 2. Results

### 2.1 Activity-Induced Rapid Primary Response Genes are Topologically Influential Across Genetic Networks in Mouse and Human Brain

We set out to investigate how ARGs interact within various types of genetic networks. Taking into account the tissue and stimulus specificity in ARGs response^12,20^, we focused on the work of Tyssowski et al.^1^ describing a set of ARGs expressed in the mouse brain as a response to brief or sustained neuronal activity (Table S1). The responsive genes were classified into three sets, rapid and delayed PRGs (rPRG and dPRG) and a set of secondary response genes (SRGs) which are activated in a temporally ordered manner^1^, as illustrated in Figure 1A. We investigated whether the three subsets of ARGs have topologically distinct characteristics by building different types of networks describing their biological interaction. To ensure the translatability of our findings in mouse to human, we ran the network analysis for both species using the original ARG set from Tyssowski et al.^1^ in Mus Musculus (Mm) and its mapped gene list in Homo Sapiens (HS). We constructed three different networks for each of the two species, namely two undirected protein-protein interaction (PPI) networks (the S-PPI network sourced from STRING database^21^ and GM-PPI sourced from GeneMANIA database^22^) and one directed regulatory network (R-NET sourced from the *RegNetwork*^23^ database). We assessed the topological influence of the different ARG-annotated groups with a set of network measures including network degree and influence (in terms of Spreading score (SS), Hubness score (HS) and integrated value of influence (IVI) score).

In the PPI networks (both in S-PPI-Hs/S-PPI-Mm and GM-PPI-Hs/GM-PPI-Mm) rapid PRGs had a significantly higher degree and influence (measured by SS, HS, and IVI scores) compared to delayed PRGs and SRGs. Since not all distributions passed the Shapiro-Wilk normality test, a Kruskal-Wallis test was performed to compare the effect of the ARG subgroup on their PPI network centralities revealing that there was a statistically significant difference in mean degree, SS, HS and IVI, between at least two groups whereas pairwise Wilcoxon test found that the mean degree, SS, HS and IVI of ARG subgroups was significantly different between rPRGs and dPRGs and between rPRGs and SRGs. There was no statistically significant difference between the mean centralities for dPRGs and SRGs, as illustrated in Figure 2. These findings were consistent for both species (Mm, Hs), as illustrated in Figure 2A-D.

**Figure 2:**
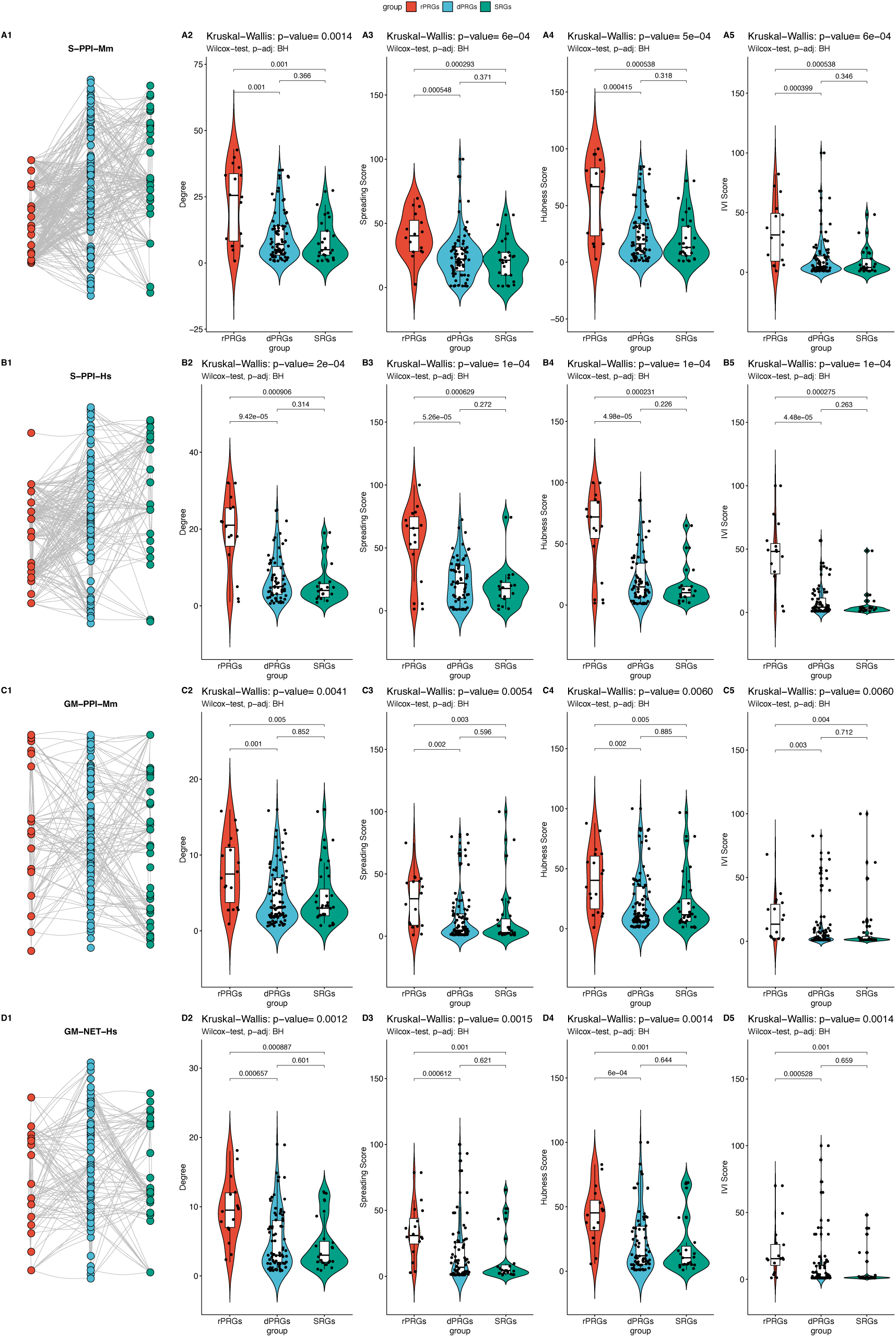
Network analysis of the undirected ARGs networks in mouse and human. Statistical comparison for centralities (column 2: degree, column 3: spreading score, column 4: hubness score, column 5: IVI score) with pairwise Wilcoxon test and Kruskal-Wallis test for the networks depicted in column 1. Rows: **A.** S-PPI-Mm **B.** S-PPI-Hs **C.** GM-PPI-Mm **D.** GM-PPI-Hs.

In the regulatory networks (R-NET-Hs, R-NET-Mm), as not all distributions passed the Shapiro-Wilk normality test, a Kruskal-Wallis test was performed to compare the effect of the ARG subgroup on its total, outgoing and incoming degree and IVI, revealing that for both species there was a statistically significant difference in mean degree and between at least two groups for the outgoing degree and outgoing IVI, while the pairwise Wilcoxon test showed that the mean outgoing degree and the outgoing IVI value of the ARG subgroup were significantly different between rPRGs and dPRGs and between rPRGs and SRGs but not between dPRGs and SRGs, as illustrated in Figure 3. Interestingly, the two R-NETs resemble a three-layer feedforward neural network with very few feedback connections from layer 2 (dPRGs) to layer 1 (rPRGs) and no feedback from the third (SRGs) group (Figures 3A1 and 3A2).

**Figure 3:**
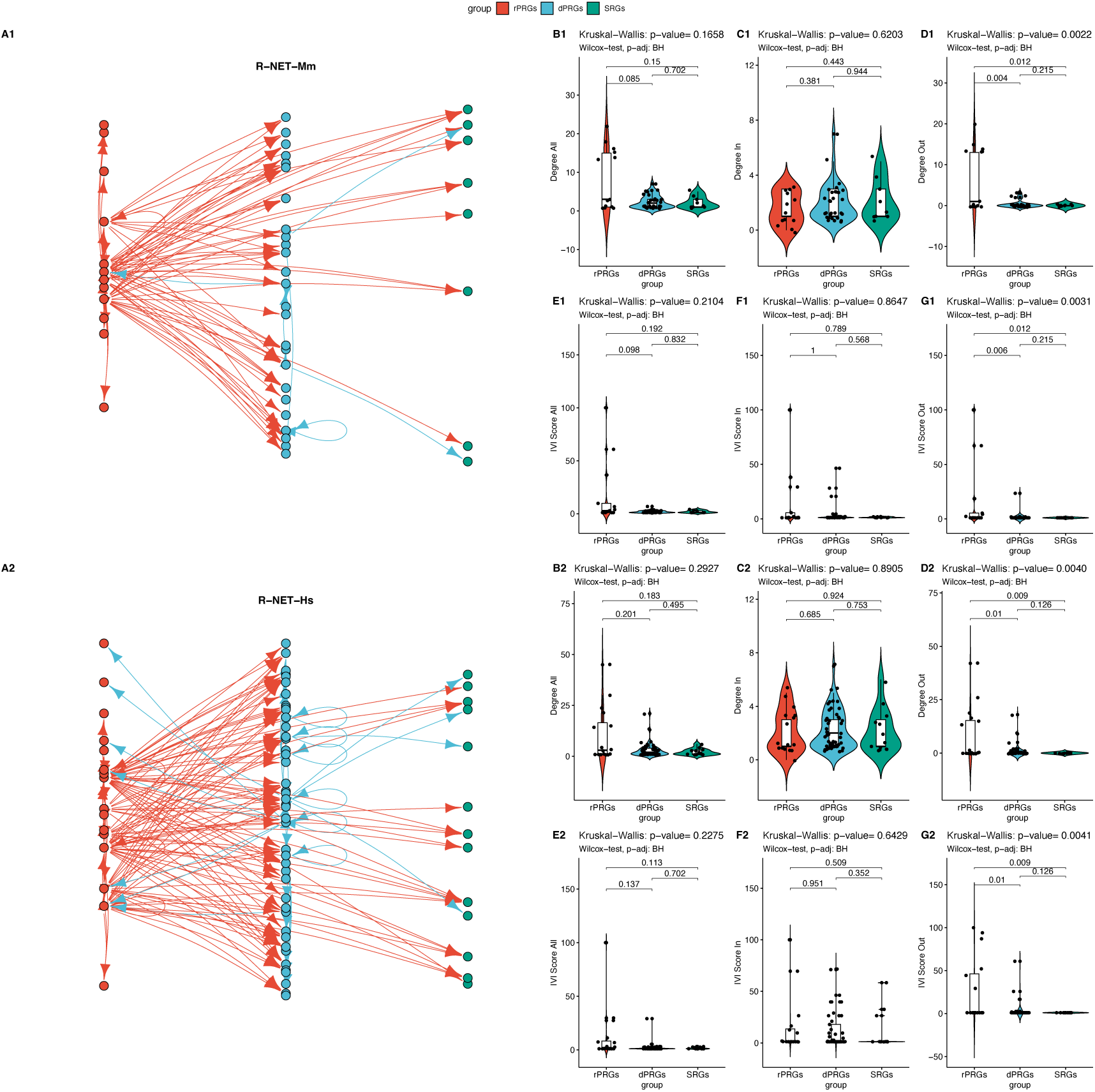
Network Analysis of the Regulatory Network. **A1-A2** Regulatory subnetworks in Mouse (top panel – R-NET-mm) and Human (bottom panel – R-Net-Hs). Comparison for the total (All), incoming (In) and outgoing (Out) degree (B1-D1, B2-D2) and IVI (E1-G1, E2-G2) centralities with pairwise Wilcoxon test and Kruskal-Wallis test.

Thus, in the context of information transfer in a directed network (based on the degree and IVI centralities), we found that in the regulatory networks, rPRGs act as transmitters to the other two layers. We also tested whether rPRGs act as sensors/receivers in the extended regulatory network including all first-degree neighbouring nodes to the core ARGs set (eR-NET-Mm, eR-NET-Hs). We found that indeed the rPRGs both receive and transmit significantly more input on average than the other two groups in both species with overall, incoming, and outgoing degree being significantly different, as illustrated in Figure 4.

**Figure 4:**
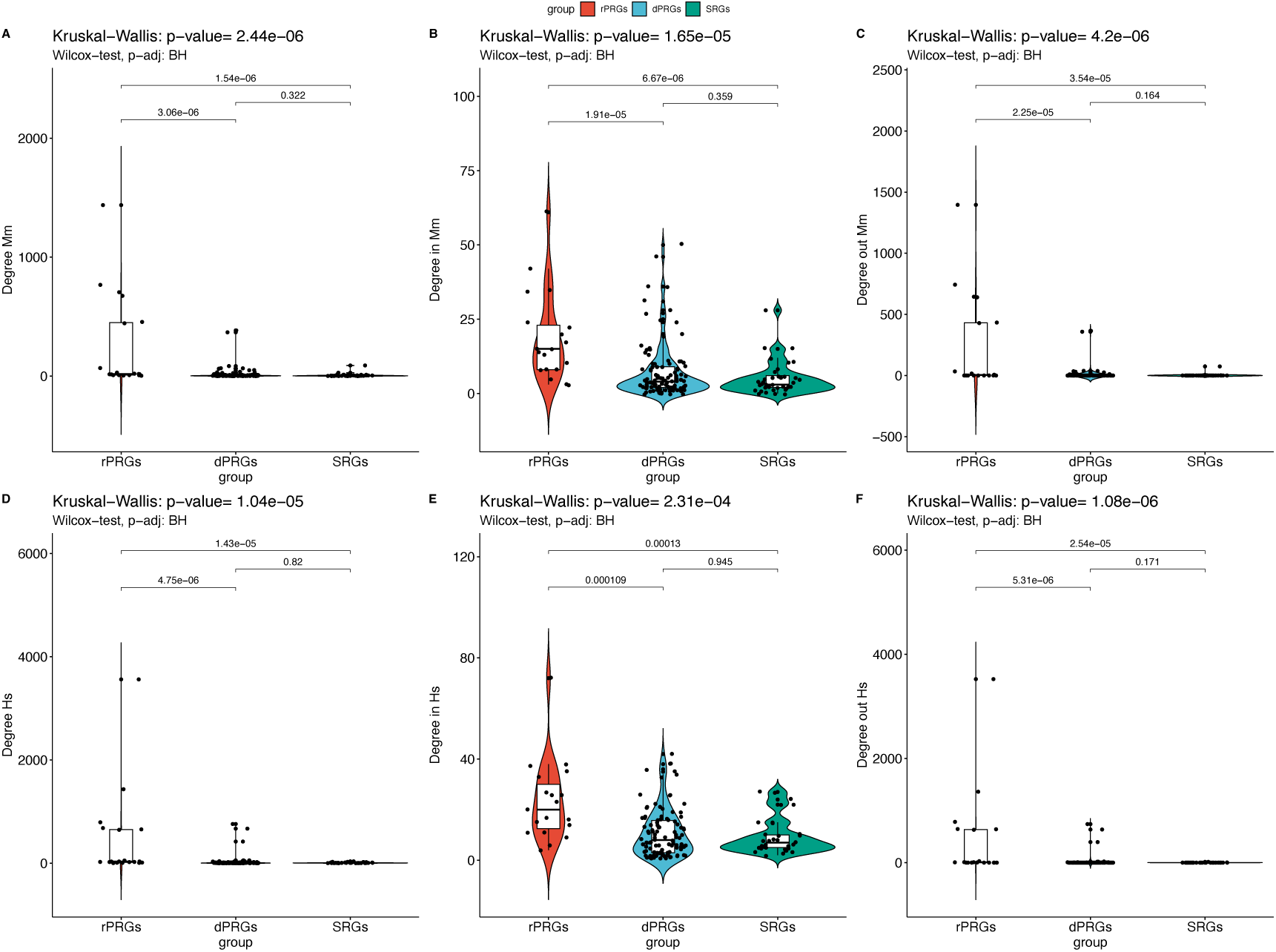
Expanded regulatory network. Top panel Mouse, bottom panel Human Comparison of total, incoming and outgoing degree distributions for ARG subgroups within the extended regulatory networks (eR-NET-Mm and eR-NET-Hs) with the first-degree neighbours with pairwise Wilcoxon test and Kruskal-Wallis test.

#### Functional Analysis

Furthermore, we explored whether the difference in topological properties in genetic networks in the three groups of ARGs also translates into dissimilar functional enrichment profiles. We performed enrichment analysis for Gene Ontology (GO) of the three ARGs groups and found that rPRGs, dPRGs and SRGs were involved in predominately different GO terms in human in mouse, as illustrated in the dot plots (Figure 5A, 5C) and the GO term-gene bipartite networks (Figures S1, S2) for the top 10 scored GO terms per group. Despite shared pathways between the two species, certain differences were observed (Tables S2a, S2b and Figure S3) with overall more enriched terms for Mm compared to Hs. The enriched terms were hierarchically clustered in functional families that are consistently found in both species in top-ranked positions (Figure 5C, 5D) such as biological processes related to skeletal muscle and memory terms for rPRGs, and blood regulation and neuropeptide signalling for dPRGs and potassium channel activity related terms for SRGs.

**Figure 5:**
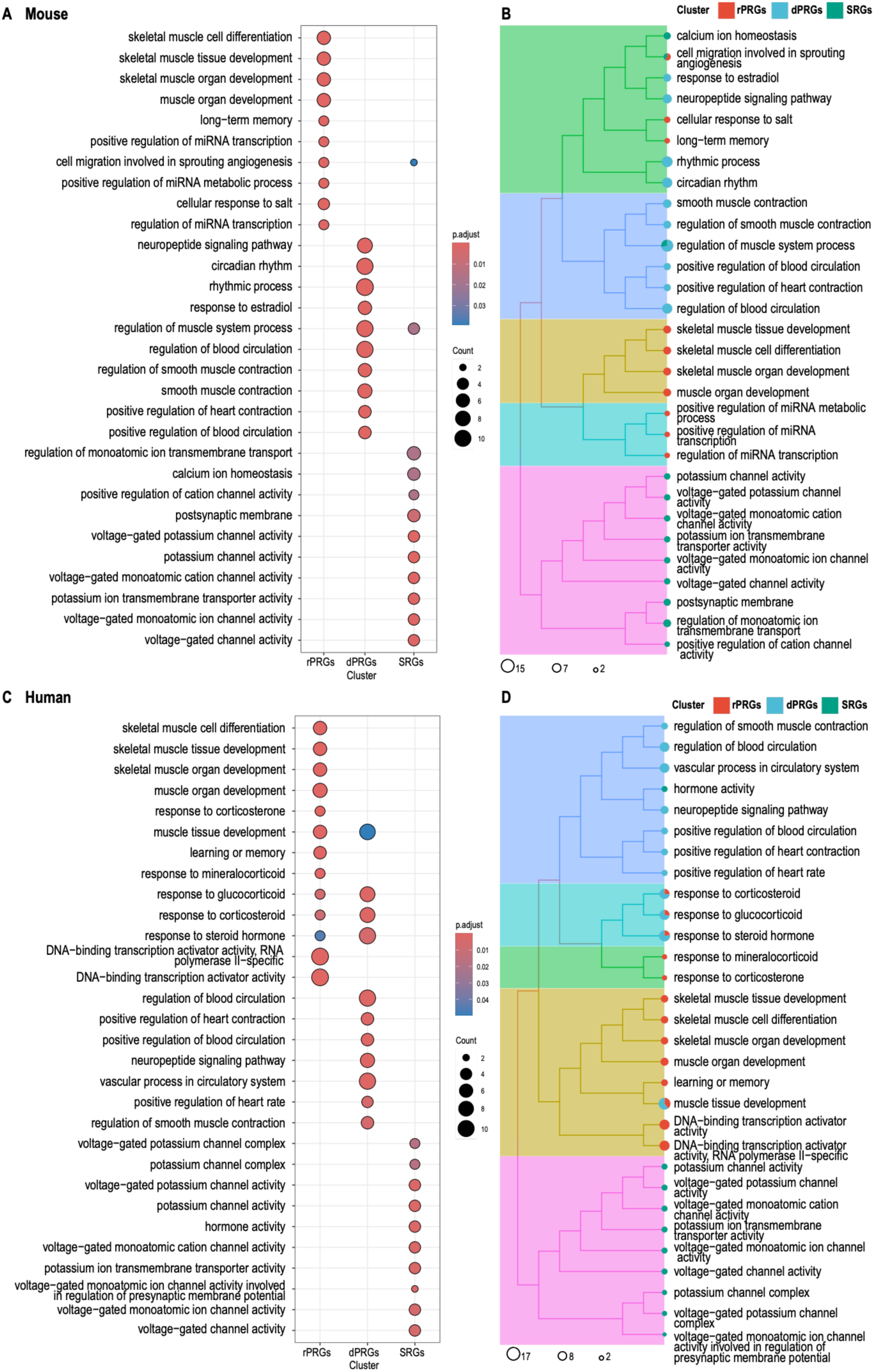
Pathway enrichment analysis. **Top Panel:** Mouse. **Lower Panel:** Human. Top 10 significantly enriched GO terms (BP, CC, MF) per ARG cluster based. **A, C:** The dot plot of enriched terms for the three ARG subsets indicates the number of genes per GO term for each gene cluster, and the colour indicates the level of the adjusted p-value. **B, D:** Heatmap of hierarchical clustering of enriched terms for the three clusters. Circle sizes indicate gene number per term.

Overall, our network analysis indicates that rapid PRGs are topologically more privileged compared to delayed PRGs and SRGs in terms of information propagation (both as sensors and as transmitters) across various types of biological/genetic networks (undirected and directed) and for both species tested here (mouse and human) and are also functionally enriched predominantly in distinct GO terms.

### 2.2 Activity regulated Genes are more genetically constrained with sparse gene-disease relevance

Following the identification of ARGs as key nodes in the flow of information in both genetic (undirected and directed) networks and the establishment of translation of these findings from mouse set to human networks, we set out to explore their role in human diseases, with an interest on brain-related diseases based on their critical role in neuronal function. We explored the disease association of our list of genes by querying DisGeNET^24,25^ for both individual diseases and disease ontologies. As illustrated in Figures 6A and 6C, there was a sparse involvement of ARGs in diseases and disease classes, respectively. The enriched disease terms and ontologies can be clustered into families that are consistently found in both species (Figure 6B, 6D). The main group categories of brain disease ontologies found were related to bipolar mood disorder and nervous system neoplasm.

**Figure 6:**
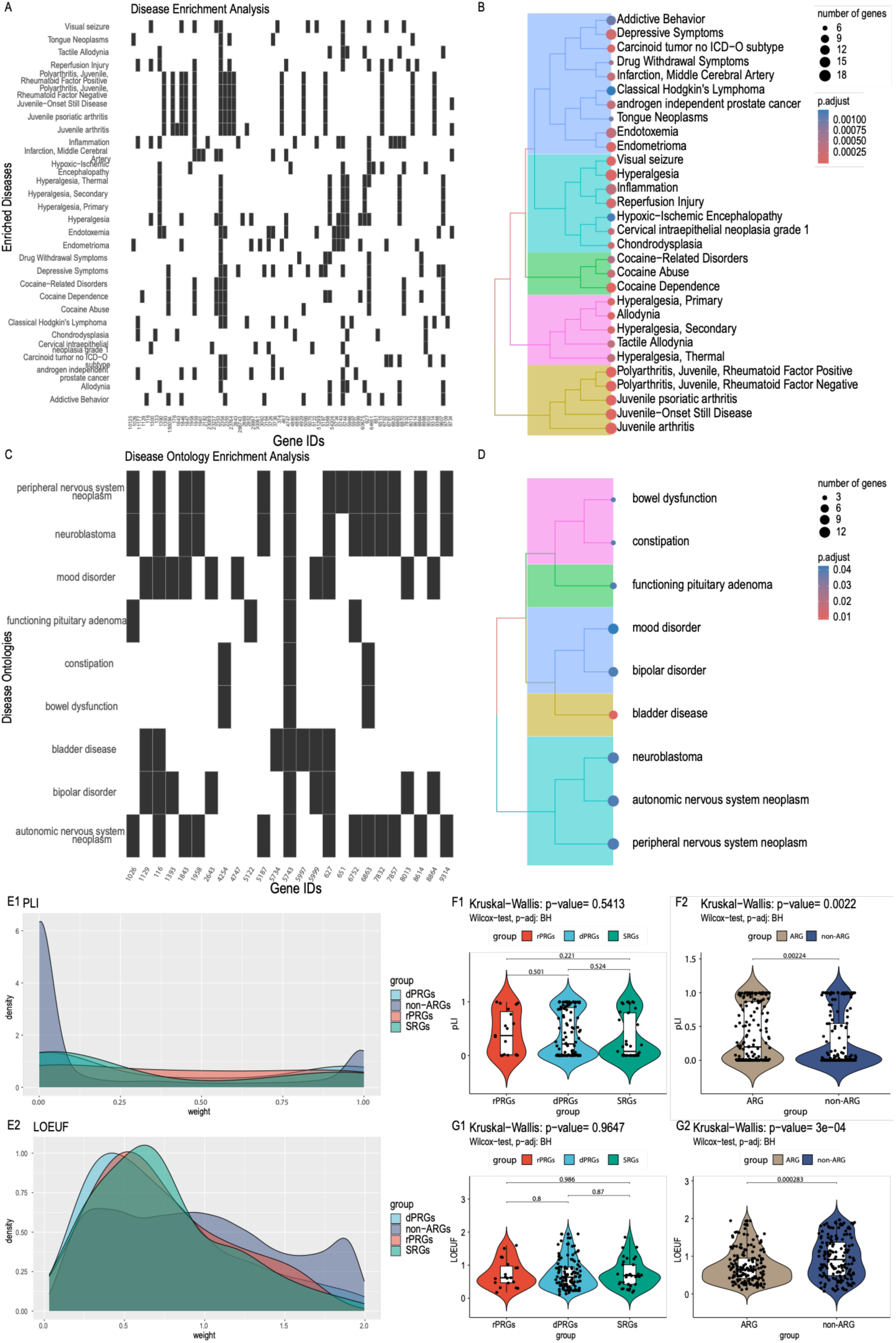
Disease load and genetic constraint of ARGs. **A-B:** Enrichment analysis results for disease and disease ontology (based on DisGeNet^24^). **C-D:** Heatmap of hierarchical clustering of the enriched disease and disease ontology terms for the three clusters. Circle sizes indicate gene number per term. **E.** Visualisation of the distributions of the genetic constraint metric pLI and LOEUF between the ARGs clusters and the full gene set available in gnomAD. **F.** Comparison of mean pLI and LOEUF among three ARG subgroups **G.** Illustrative example of comparison of mean pLI and LOEUF of ARGs with a random equally sized gene set of non-ARGs.

An explanation for the lack of a prominent association between ARGs and brain diseases could be the fact that ARGs are less tolerant to genetic variation due to high mutational constraint. To explore this possibility, we investigated whether ARGs are highly constrained and assessed that using two measures of mutational/genetic constraint metrics from the Genome Aggregation Database (gnomAD) ^26,27^, the probability of loss of function intolerance (pLI)^28^ and the loss-of-function observed/expected upper bound fraction (LOEUF)^26,27^. High LOEUF and low pLI (<=0.1) indicate loss-of-function (LoF)-tolerant genes, while low LOEF and high pLI (>=0.9) indicate LoF -intolerant genes. No significant differences were observed among the three sets of ARGs in terms of either pLI or LOEUF (Figure 6F1-6F2). However, the set of ARGs was on average more constrained for both metrics (higher pLI and lower LOEUF) compared to random sets of non-ARGs of the same gene set size (Table S3) as illustrated for one instance of a random set of genes in Figure 6G1-6G2.

Taken together, the ARG-disease analysis showed that ARGs are on average more constrained and thus less tolerant to genetic variation than a random group of genes and this could justify in part their sparser association with (brain) diseases.

### 2.3 The Role of ARGs in Alzheimer’s Disease: A Case Study

#### GWAS highlighted AD-associated genes in ARGs gene regions, mainly near FOSB

Considering ARGs’ key role in brain function, and the emerging topological influence of IEGs in genetic networks, we focused on investigating the associations between ARGs/ARG-associated variants and AD to explore whether ARGs could be directly or indirectly involved in its pathology. GWAS studies of AD have highlighted variants within several genes to be associated with disease severity^29^. Therefore, we utilized the summary statistics from a recent large-scale study on AD, which combined GWAS data from three separate cohorts, focusing on samples of European descent^29^. We identified genetic variants strongly associated with AD (*p*-value ≤ 1e-5) within a ± 200kb ARGs window (ARGs used are listed in Table S1).

From the GWAS analysis, we found 94 AD-associated single nucleotide polymorphisms (SNPs) within the expanded ARGs regions, located on chromosomes 2, 5, 15 and 19. Out of these, 73 were unique as some of the SNPs were also linked to neighbouring genes to their mapping gene after ARGs region expansion (Table S4). Only one of the identified SNPs (rs888747) was found to be located at an ARG (*CYSTM1*) whereas the rest of the 72 SNPs were located in neighbouring genes to ARGs (near *FOSB*, *EGR1*, *CYSTM1*, *COQ10B* and *RASGRP1*), as illustrated in Figure 7A. Interestingly, 58 out of the 73 variants were found to be located in 10 unique genes near *FOSB* (*PPP1R13L, ERCC1, KLC3, CKM, MARK4, OPA3, ERCC2, CD3EAP, GIPR, VASP*), none of which belonged to the initial ARG list. The effects (odds ratios (OR)) of the 73 AD-associated variants in the ARG gene regions indicated that 13 of them had a protective effect (OR < 0.95), 17 had a risk effect (OR > 1.05) and 43 of them were considered of borderline effect (0.95 ≤ OR ≤ 1.05) for AD (listed in Table S4). The effect class of variants per gene is illustrated in Figure 7B.

**Figure 7:**
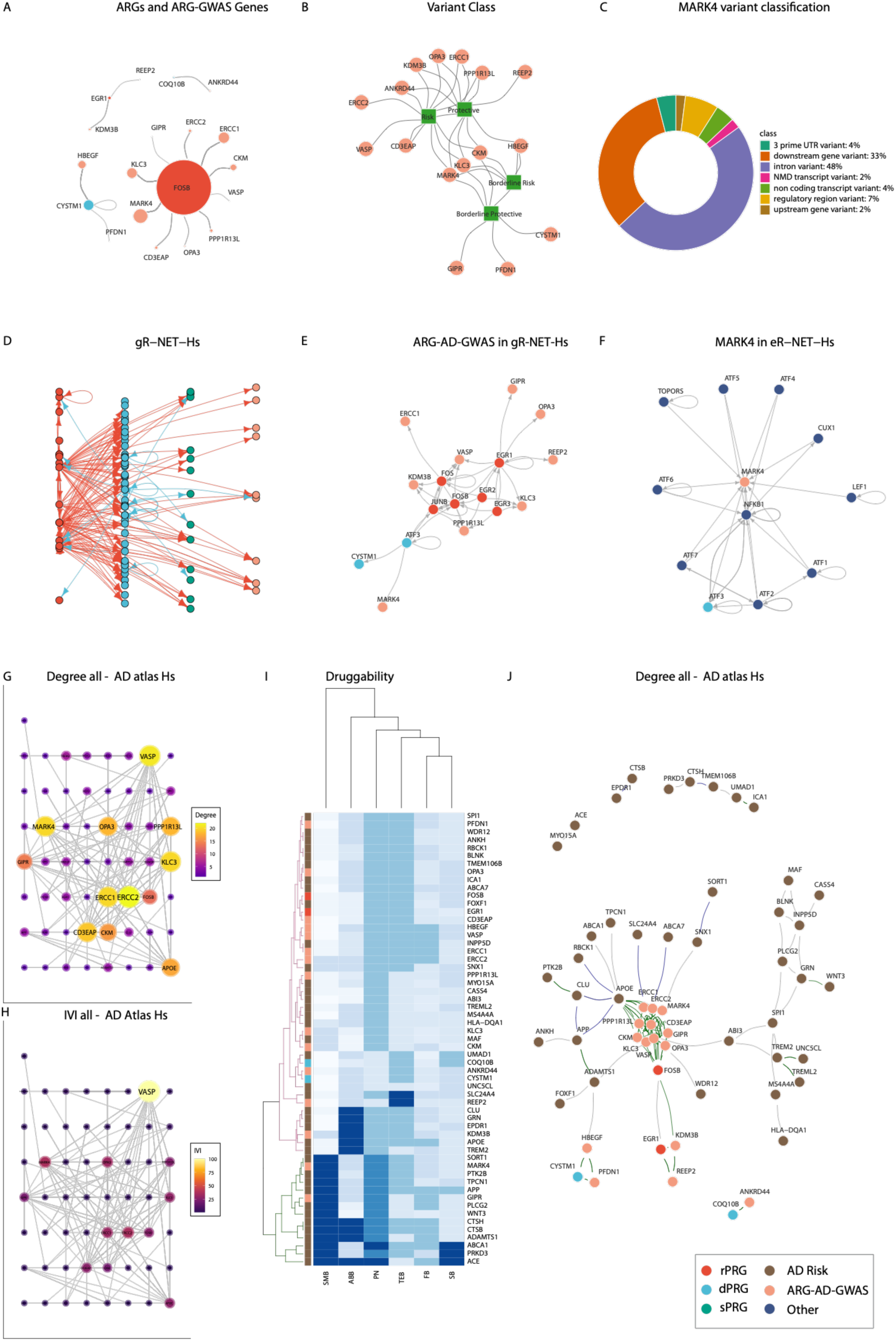
ARG-AD-GWAS genes and topological relationship to ARG in human networks and druggability. **A.** Network depiction of ARGs-GWAS Genes linked to the ARGs whose gene region they were found in. **B.** Network representation of the variant class (square nodes) of each GWAS analysis annotated gene (circle nodes) **C.** Functional annotation of the MARK4 AD-associated variants. **D.** Extended R-NET-Hs including ARG-AD-GWAS genes (gR-NET-Hs). **E.** Subnetwork of (D) with ARG-AD-GWAS and first neighbours. **F.** *MARK4* R-NET and first-degree neighbours. Co-regulation edges (dark green) co-abundance (dark blue) and co-expression (dark grey). **G-H** networks indicate different metrics or scores with size and colour range indicating the level of the respective value for the AD Atlas Network with **G.** depicting Degree and **H.** depicting IVI**. I.** Heatmap and hierarchical clustering of Agora druggability scores per gene in the AD Atlas network. The green and magenta trees indicate the higher and lower-scoring groups respectively. **J.** AD Atlas network ARG-AD-GWAS genes in beige, risk genes in brown.

##### LD analysis of ARG-AD variants

Furthermore, we ran a linkage disequilibrium (LD) analysis using the LDlink tool ^30^ to explore the LD patterns between ARG-AD risk^31^ and ARG-AD protective variants ^32^ themselves, as well as with variants in genes known to be increasing or lowering the risk of AD respectively (Table S5). LD patterns were also evaluated between the ARG borderline protective and borderline risk variants (Table S6). In addition, we wished to classify the ARG-AD borderline variants into either the risk or protective ARG groups based on LD. Results from the *LD analysis* showed that 15 borderline protective ARG-AD SNPs (0.95 ≤ OR ≤ 1.00) were in high LD (LD ≥ 0.80) with 13 borderline risk ARG-AD SNPs (1.00 < OR ≤ 1.05). None of these variants were in high LD with AD-GWAS SNPs with either protective or risk effects. The variants are located on chromosomes 5 and 19, spanning the gene regions of *HBEGF, MARK4, CKM, KLC3* and *GIPR* with most of them located in *MARK4*. Furthermore, none of the borderline protective ARG-AD SNPs were in high LD with the protective ARG-AD SNPs or any other of the AD-GWAS SNPs with protective effects on corresponding chromosomes/regions. On the contrary, four borderline risk ARG-AD SNPs (rs11666411,rs11672894,rs11672923,rs11667235) (LD = 1) were in high LD with a risk ARG-AD SNP (rs344812) (LD > 0.9) but not with any risk AD-GWAS SNPs in the corresponding regions. At the same time, the risk ARG-AD SNP rs344812 was also in high LD with four borderline protective variants (LD ≥ 0.94). The ARG-AD variants found to be in high LD were all in the *MARK4* gene region. This, in turn, indicates the uncertainty of the effect of these borderline variants on AD which require further investigation.

#### The ARG-GWAS analysis highlights *the role of* MARK4 in Alzheimer’s disease

The ARG-AD-GWAS analysis highlighted the *FOSB* gene region as the region with the majority of AD-associated variants (*p* ≤ 1E-5). The ARG-AD-GWAS variants were located in 10 unique genes (Table S4) near *FOSB* while *MARK4* was in the spotlight due to containing the vast majority of the detected AD-associated SNPs.

Thus, we focused on *MARK4* variants included in our ARG-AD-GWAS list and performed *MARK4* variant classification and functional annotation. We classified the *MARK4* variants as previously mentioned (see Table S4 and Methods); two of them had a protective effect, one of them had a risk effect and another fifteen had a borderline effect. We aimed to further investigate these to potentially observe previously unseen AD mechanisms.

To facilitate the functional investigation of *MARK4* variants with protective, borderline or risk effect, we used data from the Accelerating Medicines Partnership for Alzheimer’s Disease (AMP-AD) Consortium ^33^ to explore the association of these variants with gene expression through the expression quantitative trait loci (eQTLs) associated with *MARK4*. As we had already explored the LD patterns between the *MARK4* variants of our ARG-AD list and the variants in either AD protective or high-risk genes, we examined which of these variants were also eQTLs with a particular focus on the rs344812 risk SNP, which was in high LD with additional borderline *MARK4* variants. Both the borderline protective and borderline risk *MARK4* variants were also in high LD between them (Table S7).

##### MARK4 eQTLs in the brain

Moreover, we investigated whether the eQTLs of the ARG-AD list were expressed in the brain, specifically in the cerebellum (CER) region from the Mayo study and in cortical regions from a meta-analysis study including four cohorts (ROSMAP, Mayo, MSSM-Penn-Pitt and HBCC)^33^. The cis-eQTL list of the RNAseq cohorts (AMP-AD) showed that all *MARK4* variants except the two protective ones (rs149628060, rs183442275) were eQTLs. These protective SNPs are both rare (Minor Allele Frequency (MAF) < 5%) and no studies have previously reported them. The results of the cerebellar regions did not show statistically significant associations of the *MARK4* genetic variants with gene expression. On the other hand, the analysis using the cortical regions indicated that rs344812-A, which was found to increase the risk of AD in the ARG-GWAS analysis, also showed increased expression of *MARK4* (OR = 1.14, p = 1.8 x 10^-3^, FDR = 0.06).

Furthermore, we searched for the functional consequence of each of the eQTLs using the Variant Effect Predictor (VEP) database^34^. The *MARK4* variant functional annotation results from VEP are shown in Figure 7C. The majority of the SNPs were introns and downstream gene variants while 7% were regulatory region variants and 4% non-coding. The risk variant rs344812 in particular was also an intron while the protective variants rs149628060 and rs183442275 were an intron and a 3’-UTR respectively.

Taken together, the GWAS analysis here highlights the potential impact of the *FOSB-* neighbouring region in AD and provides suggestive evidence of *MARK4* as one of the driver genes contributing to AD pathogenicity.

### 2.4 Network analysis of ARG-AD-GWAS network neighbourhood highlights potential therapeutic avenues for AD

#### MARK4 and other ARG-AD-GWAS genes are downstream regulated by PRGs

The ARG-AD-GWAS analysis highlighted key genes associated with AD in the ARGs’ genomic regions; all ARG-AD-GWAS were found close (in terms of chromosome location) to 5 ARGs; 2 rPRGs (*FOSB* and *EGR1*) and 3 dPRGs (*CYSTM1*, *COQ10B* and *RASGRP1*), as illustrated in Figure 7A. We then set out to explore how ARG-AD-GWAS are placed in terms of topology in genetic networks. Firstly, we explored whether ARGs serve as upstream or downstream regulators of the ARG-AD-GWAS genes. To this end, we revisited and expanded the ARG regulatory network R-NET-Hs to gR-NET-Hs to include the ARG-AD-GWAS genes (Table S4). We observed that several ARG-AD-GWAS genes (*KLC3*, *GIPR, REEP2*, *VASP*, *OPA3*, *PPP1R13L*, *ERCC1*, *KDM3B*) are predominantly regulated by a set of rPRGs including *FOSB* as well as *EGR1*, *EGR2*, *EGR3*, *FOS*, and *JUNB* and the dPRG *ATF3* (Figures 7D-E). These ARGs represent the top regulators (top 90 for degree and IVI) Table S8) both in the gR-NET-Hs network (Table S8) and in the original R-NET-Hs network (Table S9) and are all direct regulators of transcription (Table S1).

Focusing on *MARK4* (Figure 7E), its only regulator was *ATF3,* which also regulates *CYSTIM1* (the only ARG that had an AD-related SNP albeit with borderline OR). We then investigated all contacts (including from non-ARGS) of *MARK4* in the expanded R-NET subnetwork (with ARGs and their first neighbours) and found that MARK4 is regulated by the whole ATF gene family and additionally by the *TOPORS, CUX1*, *LEF1* and *NFKB1 genes* (Figure 7F).

Taken together, this analysis shows that the ARG-AD-GWAS genes are downstream regulated by PRGs and highlights the potential role of PRGs in the regulation of AD-associated genes.

#### MARK4 in a closely linked cluster with ARG-AD-GWAS in AD multi-omic network

We further explored the topological characteristics of the ARG-AD-GWAS genes in the Alzheimer’s Disease Atlas (AD Atlas)^35^, an integrated multi-omic network resource around AD, using as input the combined set of the AD risk genes (Table S5) including *APOE*, the ARG-AD-GWAS genes and the ARGs in whose neighbourhood the GWAS genes were found in (Figure 7D). In this multisource AD atlas network (filtered with brain region), we found that ARG-AD-GWAS were closely connected/coregulated (eQTL) among them and with the ARGs in whose genomic region (or close to) their SNPS were found in. The most tightly linked module was the set of genes proximal to *FOSB* (Figure 7J). This dense *FOSB* cluster was also closely linked to *APOE*. This modular structure was found on a smaller scale for the other ARGs and their nearby genes (around *EGR1* and *CYSTM1*) and these two smaller clusters were also connected to the *FOSB* main module. This modular structure was not observed among known risk genes for AD.

Through network analysis of the AD Atlas network, we found that *MARK4* was consistently ranked highly (top 95% quantile) both in terms of degree and IVI (see centralities in Table S10 and Figures 7G and 7H). Other genes of high topological importance in terms of degree/influence included the risk genes *VASP* and *ERCC2*, two genes that were notably linked to AD only with risk SNPs (Figure 7B). Other than the AD-ARG-GWAS genes, *APOE* and *FOSB* were also highly ranked topologically. This network analysis also showed interesting communication paths between the ARG-GWAS-AD clusters linked with ARGs and AD risk genes.

Taken together, this analysis highlights the intricate relationship of ARGs and ARG-AD-GWAS a tightly coupled module of interest with known risk AD genes to show less modularity and being allocated in the periphery of these clusters.

#### MARK4 among the key AD druggable targets near ARGs

Following the topological analysis, we focused on exploring the druggability potential of the highlighted modules and other annotated risk genes in the AD Atlas network. Toward that end, we retrieved from AMP-AD the complete Agora list of AD drug targets that have been nominated by AD researchers. The AMP-AD pharmaceutical partners have scored several drugs of interest to AD for their druggability/ligandability based on a multistep process ^36–41^. Here we retrieved and normalised the scores (assigned in buckets) in [0,1] (with 1.0 indicating the optimal score) for further comparison for the following classifications: small-molecule drug development (SMB), therapeutic antibody feasibility (FB), safety_(SB), ABability (ABB), new modality (NMB), tissue engagement (TEB) and Pharos classification number (PN). We then classified genes using hierarchical clustering based on these druggability features. As illustrated in Figure 7I (green part of the hierarchical tree), the top in terms of overall druggability scores out of the two main clusters included several AD risk genes and two AD-ARG-GWAS genes: *MARK4* and *GIPR*.

Overall, exploring the topological features of the ARG-AD-GWAS highlighted genes in the extended gene regions of ARGs (focus on ARG and GWAS friends *FOSB* dense neighbourhood) pinpointed key neighbouring targets amenable to pharmacological/therapeutic intervention in AD, with the spotlight on *MARK4*.

## 3. Discussion

IEGs have been extensively studied as indirect markers of neuronal activity and as specific genes and the type of stimuli that elicit their response ^42^. Our work here has expanded our knowledge towards two main directions by delving into the influential role of IEGs in genetic networks and exploring the key links of IEGs, ARGs and nearby genes to AD to shed light on potential therapeutic avenues for further investigation.

### IEGs serve as influencers across different types of genetic networks in mouse and human

Here we have pinpointed the rPRGs/IEGs characteristics in being influential members of ARGs genetic networks. These findings are conserved not only across different types of networks (undirected: PPI and integrated PPI and directed: regulatory), but also across two species, human and mouse. Thus, ARGs’ subclasses are distinct not only in terms of the timescale and mode of activation (as extensively discussed in ^5^) but also in terms of their topological influence in genetic networks. This finding is also consistent at the level of functional level as suggested from our analysis in which we found rapid PRGs, delayed PRGs and SRGs to be involved in predominately different GO terms.

Notably, rPRGs are topologically more privileged compared to the other two groups in terms of information propagation both as sensors and transmitters. The influential role of rPRGs as sensors is consistent with their role as early responders to stimuli and as transmitters with the dual role of some rRPGs as transcription factors ^5^. The combination of the two features poses a strategic advantage for rPRGs to be able to receive and pass down a transcriptional wave to downstream targets, much like an input layer in a feedforward neural network.

### ARGs are sparsely involved in diseases likely due to their high mutational constraint

A key finding of this work was its translational aspect. The list of ARGs recorded in mouse was used to construct genetic networks and the analysis revealed the same topological properties in both mouse networks and human homolog networks, across all tested scenarios, further strengthening our investigation for IEGs for human diseases.

The strategic topological position of IEGs within the ARGs network motivated us to consider them in terms of their relevance to brain diseases. In addition, we explored the possible role of all ARGs in disease pathogenicity. We found that ARGs were sparsely involved in diseases, including AD, from gene-disease databases. We hypothesized that this could be attributed to the fact that ARGs, including rPRGs, are also less tolerant to genetic variation. Our analysis showed on average a significantly higher mutational constraint for ARGs versus non-ARGs. These results are consistent with work showing that, compared with a random group of genes, highly constrained genes (based on pLI) are significantly more likely to be associated with disease of the Mendelian type^43^. Although highly constrained genes are intolerant to mutational variation this does not disqualify them from serving as possible drug targets. Even essential genes that are intolerant to loss-of-function variants can still be successfully targeted with inhibitory drugs ^44^. Whether ARGs would serve as optimal therapeutic targets, given their widespread influence in genetic networks despite their intolerance to mutations remains to be determined.

### Genetic variants (SNPs) significantly associated with AD in genes near ARGs: Focus on FOSB and MARK4

With AD as a case study, using GWAS summary statistics, we found only one SNP within an ARG region while all other SNPs were located in neighbouring-to-ARG gene regions. Most of the SNPs significantly associated with AD were found in ten unique genes near a well-known IEG, *FOSB*. Although *FOSB* has not been linked to AD, its truncated splice variant *ΔFosB* (with an atypical much longer half-life of days compared to hours for *IEGs*) was shown to regulate gene expression and cognitive dysfunction in an AD mouse model ^45^. An interesting finding here was that several SNPs showed a borderline effect on AD that led to uncertainty in classifying them into the protective or risk groups, as the LD analysis only showed relationships between themselves and not with other known risk or protective variants. Most of these variants were in *MARK4*. Notably, an AD-risk variant was also found to be an eQTL associated with an increased expression of *MARK4* in cortical regions but not in the cerebellum. This is an intron variant, but further investigation is needed to establish the consequences of the eQTL on AD pathogenicity.

### MARK4 and other ARG-AD-GWAS genes are downstream regulated by PRGs

Our analysis within the regulatory network showed that the ARG-AD-GWAS genes are downstream regulated by key PRGs in terms of topological influence and highlights the potential role of PRGs in the regulation of AD-associated genes. This contributes to the knowledge gap in mapping upstream regulators and/or downstream targets of ARGs^12^ and particularly PRGs. Some of the genes such as *Egr1* and *Fos/c-Fos* (*Bdnf* - dPRG and Arc - rPRG) have also been linked with AD through the downregulation of their expression by *APP* in the prefrontal cortex of mice^15^.

Among the non-ARG genes targeting *MARK4* were the ATF family of genes and *NFKB1.* ATFs have been suggested as potential causative genes and drug targets for AD due to their significantly altered expression in AD^46^. *NFKB1* is also a gene of interest in AD due to its role in brain inflammation and neurodegeneration as a transcription factor and master regulator of genes key for various processes including inflammation, cell cycle, proliferation and death^47^. Thus, the *MARK4* gene is also highlighted as the target of non-ARG genes important for AD.

### MARK4 a druggable AD target in dense AD multi-omic cluster of ARG-AD-GWAS

*MARK4* encodes for MAP/microtubule affinity-regulating kinase 4 and is involved in microtubule organization in neuronal cells. The expression of *MARK4* is elevated in the brains of AD patients and its activity colocalizes with early pathological changes ^48^ (through pathological phosphorylation of tau protein). Also in a fly model, Oba et al reported that MARK4^ΔG316E317D^ increases the abundance of highly phosphorylated, insoluble tau species and exacerbates neurodegeneration.

*MARK4* has recently been in the spotlight due to its promising potential as a drug target ^49–51^. Specifically, Shamsi et. al reported that the inhibition of *MARK4* by serotonin can act as a therapeutic approach to tackle AD and neuroinflammation ^51^. In our recent computational work, the serotonergic synapse pathway was highlighted as top scoring in a weight-modulated majority voting of the modes of action (MoAs), initial indications and targeted pathways of the drugs in clinical trials and second in proposed from computational repurposing studies in AD ^52^. The MoA serotonin receptor antagonist was also highly ranked in both cases ^52^.

Although *MARK4* has been previously proposed as a potential drug target for AD and has a high-ranking druggability score on the Agora list of AD drug targets, it has not been explored extensively and so far no clinical trials have focused on *MARK4* inhibitors for AD. A possible challenge for targeting *MARK4* could arise due to the multiple roles of MARKs and the need to maintain MARK activity at a physiological level for normal neural transmission ^49,53^. Thus, further studies are required to explore the implications of targeting *MARK4* directly or indirectly (through its regulators). Such indirect treatment strategies for *MARK4* could benefit from focusing on the tight cluster of genes (containing *MARK4* and the other genes with AD-associated variants in the genomic neighbourhood of *FOSB*) which has emerged from our analysis in the AD brain-specific multi-omic network in the AD Atlas network. An indirect approach that targets IEGs, could capitalise on other types of non-pharmacological interventions by harnessing the activity of IEGs rather than targeting them pharmacologically. Recent studies have shown that induction of gamma oscillations with visual stimulation results in the reduction of amyloid plaques and phosphorylated tau in various AD mouse models and also induces a set of IEGs that include the *Mark4* regulator *Atf3* ^54^. Later studies have also shown similar effects with other types of sensory stimuli like auditory ^55^ and tactile stimulation ^56^.

### Implications

Despite recent promising efforts towards developing treatments in AD^57,58^, there is an urgent need for both pharmacological and other interventions to halt/delay the onset of the disease. Our work contributes to highlighting *MARK4* as a potential drug target that has not been explored sufficiently to date. Combining the findings, following the GWAS analysis and the network analysis in AD, we highlighted *MARK4* as the most interesting potential target due to (1) containing most of the AD-associated variants (2) containing an eQTL found to elevate the risk of AD and increase the expression of *MARK4* in the cerebral cortical region (3) containing several borderline variants which deem further exploration (4) being the target of several PRGs including AD-linked genes (5) having an influential topological position in the AD multi-omic network (6) having a highly ranked druggability Agora profile for AD targets.

### Ideas, Speculation and Recommendations

Having focused on ARGs due to their influential role in networks and combining with GWAS analysis, we have explored in detail the genomic area of *FOSB* and highlight *MARK4* as a promising underexplored drug target through additional network analysis, which we might not have focused otherwise. We speculate that implementing this approach in other diseases could serve as a good strategy for searching for possible disease-modifying targets and direct/indirect ways for pharmacological interventions by mapping and analysing their upstream and downstream neighbourhood, i.e., ‘friends’ of influencers such as IEGs.

### Limitations

One limitation of this work is that it potentially does not include all IEGs/ARGs in the brain as ARGs are elicited in response to different types of stimuli. Another limitation is that there are discrepancies in the categorization of these ARGs in terms of rapid or delayed activation. This is consistent with what has been reported in Tyssowski et al. ^5^ of certain genes being classified differently depending on the type of the experiment (in-vivo or in-vitro). Also, Homer1a which has been highlighted as a key marker and therapeutic target for AD was not considered in our analysis as it was excluded (along with *Vgf, Gadd45b* and *Nfkbid*) from the ARG list generated in Tyssowski et al. due to ambiguity on its classification based on induction kinetics ^5^. One weakness that could be attributed to the experimentally obtained list being in mice was alleviated by showing the translational aspect in humans of consistent network characteristics of IEGs in both species.

### Future Plans

Overall, our approach of combining network and GWAS analysis has provided a framework that can be extended to other brain diseases, such as neuropsychiatric and neurological diseases^59^. A key outstanding challenge is not only uncovering the neighbourhood in regulators but also the specific ARG networks to tissue and cell signalling patterns. The application of this is also extended to uncovering region- and cell-specific ARG subnetworks of interest for AD and other brain diseases.

### Conclusions

We have explored the critical role of IEGs in genetic networks and potential drug targets for AD within or close to ARGs. This contribution is important because it fills the knowledge gap on how IEGs interact at the network level, and this system-level investigation proved beneficial in studying potential targets for AD, a most prevalent brain disease. With a combination of GWAS analysis and network approaches, *MARK4* was highlighted as a highly relevant and understudied potential pharmacological target for AD. Overall, we demonstrated the added value in studying IEGs/ARGs in the network context, both for a better understanding of their communication and to pinpoint genes of clinical interest. Given the key function of IEGs as temporal signal integrators and orchestrators of gene expression programs, elucidating the influential role of IEGs in networks can provide valuable insight into developing therapeutic interventions for diseases where IEGs and their downstream targets are dysregulated.

## 4. Methods

### 4.1 ARGs List pre-processing and mapping

We mapped the mouse gene symbols as obtained from Tyssowski et al.,^5^ to their ortholog human gene symbols and for both species, we also mapped them to their EntrezIDs using a merged output from the homologene^60^, babelgene^61^ and org.Mm.eg.db^62^ R packages to maximize the number of genes that could be mapped in terms of gene symbols and EntrezIDs. In total all but three genes (*4931440P22Rik, 5430416O09Rik, Glt28d2)* were mapped successfully from Mus musculus (Mm) to Homo sapiens (Hs). The full mapped list is available in Table S1.

### 4.2 Network Construction

For the analysis in section 2.1, we constructed three different networks for each of the two species as detailed below:

#### S-PPI Network

We constructed two PPI networks for the two species (Mm and Hs) respectively for the ARGs set based on known PPIs verified by experiments from the STRING database^21^. The STRING database^21^ was downloaded (Mm and Hs Files: *10090.protein.physical.links.detailed.v11.5.txt*,*9606.protein.physical.links.detailed.v1 1.5.txt*). The R package biomaRt^63^ was used to map their ensemble ID to gene symbol, EntrezID and UNIPROT ID. We selected only the PPIs with experimental evidence with no cut-off. The graph was simplified by averaging the weights of multiple edges between two nodes. The unweighted version of the S-PPI-Mm and S-PPI-Hs was used for further analysis.

#### GM-PPI Network

We constructed two integrated networks for the ARGs set (one for each species Mm and Hs) based on the integrated known gene relations from the GeneMANIA^22^ database by querying the webserver https://genemania.org/ (Date: 15 February 2022). We selected 3 types of networks: *co-localisation*, *Physical Interactions* and *Shared protein domains*. The network weighting method was set to be *Automatically selected*. The *GeneMANIA* integrated network was constructed based on the weighted sum of the pairwise weighted edge vectors (for each pair of genes) for these three types of networks. The unweighted version of the GM-PPI network was used for further analysis.

#### R-NET Network

We constructed two regulatory networks (one for each species: Mm and Hs) based on known gene regulatory relations from the *RegNetwork*^23^ database. Data were downloaded from the repository Regulatory Network Repository (RegNetwork, https://regnetworkweb.org/, last updated 2019)^23^ for both mouse and human regulatory networks. To assess the properties of the ARGs set in section 2.1 a subnetwork was extracted containing only the ARGs list of genes in mouse and human respectively (microRNAs were excluded). We also considered the extended full network for the analysis in the Results section 2.4. The unweighted version of the network was used for further analysis.

### 4.3 Network Analysis

We used the igraph package in R (http://igraph.org/r/)^64^ to process the data and to create, analyse and visualise the networks of interest. For the networks that were not fully connected, we selected the largest fully connected subgraph component (largest diameter). The connectivity and influence within the networks for each ARG subgroup were assessed by calculating the following local and global network centralities: (1) node’s degree (using the igraph R package^64^), which quantifies the number of nodes each node of interest is connected with, (2) node’s spreading score (SS), which quantifies the spreading potential of each node, (3) the node’s hubness score (HS), which quantifies the power of a node in its environment and (4) the Integrated Value of Influence (IVI) which combines the local, semi-local and global centralities to unify them in a single influence score. Both HS and SS are major components of IVI and all three were calculated using the R package Influential ^25^. For directed networks, we calculated the centralities for all and separately for both incoming and outgoing edges.

### 4.4 Network Statistical Analysis and Visualisation

The statistical tests for the network centralities stratified based on group membership were performed in R (version 4.1.1). The different centrality distributions did not satisfy the normality Shapiro-Wilk test. Thus, we used Kruskal-Wallis (a non-parametric alternative to the one-way ANOVA test) to test the effect of ARG subgroup membership and the pairwise Wilcoxon test to compare group levels pairwise with corrections for multiple testing (using the Benjamini & Hochberg (BH) p-value adjustment method). For the statistical analysis and visualisation of the networks^64^ also the packages RColorBrewer ^65^, ggplot2 ^66^, ggpubr ^67^, rstatix ^68^, dplyr ^69^, ggsci ^70^, gplots ^71^, tidyr ^72^, scales ^73^, plyr ^74^, ggplotify ^75^ were used complementary to igraph^64^ and Influential ^25^.

### 4.5 Functional Enrichment Analysis

The overrepresentation functional enrichment analysis was performed with the R package clusterProfiler^76^ with reference database the Gene Ontology (GO). The p-value was adjusted using the Benjamini-Hochberg correction for multiple comparisons (p < 0.05). The R package ggvenn^77^ was also used to visualise the differences and commonalities for the significant GO terms for each GO partition, ARG subgroup and species for Figure S3.

### 4.6 ARG – Disease and Constraint Analysis

#### Disease Enrichment Analysis

The gene-disease association was performed by querying DisGeNET ^24^ using the clusterProfiler^76^. In addition, the packages DOSE^78^, GOSemSIM^79^ and enrichplot^80^ were used for the analysis and visualisation of the disease enrichment analysis.

#### Mutational Constraint Analysis

The Genome Aggregation Database (gnomAD) was accessed on 11/01/2023 (downloaded file: gnomad.v2.1.1.lof_metrics.by_gene.txt). We cleaned the gnomAD data file for duplicates and annotated which genes are ARGs and of which type. For the gene constraint analysis, we scored genes based on the constraint using the “loss-of-function observed/expected upper bound fraction” or “LOEUF” and the *probability of being loss-of-function intolerant* (pLI) scores available in gnomAD^26^. A total of 1000 random non-ARG gene sets were generated (of equal size to the ARG gene set). We then compared the pLI and LOEUF distributions of each of the random non-ARGs set to that of the ARG gene set mean (Kruskal-Wallis test and pairwise Wilcoxon test with the alternative hypothesis “greater” for LOEUF and “less” for pLI).

### 4.7 AD-GWAS Analysis

#### GWAS summary statistics

We used GWAS summary statistics ^29^ to identify all SNPs which were strongly associated with AD in European populations, within a ± 200kb ARGs window. We initially filtered the variants showing evidence of AD association at genome-wide significance (p ≤ 5e-8), but due to limited findings, we used a less stringent threshold (p ≤ 1e-5) to maximize the number of variants. Based on the effect size of the variants identified, we considered three groups: protective variants (OR< 0.95), borderline variants (0.95 ≤ OR ≤ 1.05) and risk variants (OR > 1.05). Borderline variants were split into borderline protective (0.95 ≤ OR ≤ 1.00) and borderline risk (1.00 < OR ≤ 1.05). The analyses were implemented in R-version 4.1.

#### Linkage disequilibrium analysis

The linkage disequilibrium (LD) analysis was performed through the LDmatrix of the LDlink tool ^30^ (https://ldlink.nci.nih.gov/?tab=home), using rsIDs as input. In line with the GWAS analysis, we only used European populations to conduct the LD calculations. An LD ≥ 0.80 was considered as high. We also used a list of ARG-AD risk (Supplementary Table 5 in ^31^) and ARG-AD protective variants (Supplementary Table 1 in ^32^) to explore the LD patterns between literature-based variants and the ARG-AD-GWAS variants. Since the region corresponding to the *APOE* gene was excluded from the list of the AD-risk genes ^31^, we used PhenoScanner ^81,82^ to identify and incorporate the variants associated with AD pathogenicity in the ARG-AD risk list (p ≤ 1e-5).

#### Functional annotation

eQTL results from the meta-analysis of the four cohorts in cortical brain regions (synapse id: syn16984815) and Mayo study in cerebellum region (synapse id: syn16984411) were retrieved from the AMP-AD Consortium to facilitate the identification of *MARK4* eQTLs expressed in brain regions ^33^. The data included the eQTL summary statistics as well as the increasing-expression allele. The eQTL analysis was performed in R-version 4.1.

#### Variant Functional consequence

The Variant Effect Predictor (VEP) database^34^ (https://asia.ensembl.org/Tools/VEP) was used to uncover the functional consequence of the *MARK4* risk, protective and borderline variants. Rs IDs were used as the input data to the VEP algorithm.

### 4.6 Post-GWAS Network Analysis

#### AD Atlas

AD Atlas is a network-based framework for AD which provides access to integrated multi-omic data for AD ^35^ and is available at https://adatlas.org/. We build the AD Atlas using as input the ARGs list, the ARG-AD-GWAS list and the risk list from ^31^, including APOE, with settings (Type: gene, Cutoff: genome-wide, Expand: no, Dist: 0). We filtered out the AD traits and non-gene elements from the network. The network included co-regulation edges (dark green) co-abundance (dark blue) and co-expression (dark grey) links and was undirected. The centrality analysis and visualisation were done in line with the previously mentioned networks.

### 4.7 Druggability Analysis

We retrieved the druggability information for our genes of interest from the AD knowledge portal ^83^. The AD-drug list file was retrieved from synapse.org (synapse id: syn13363443). The Ensembl IDs were mapped to gene symbols using the R package org.Hs.eg.db^62^. We then normalised the scores by inverting the values assigned in buckets to scale them in the (0,1] range, with 1.0 indicating the most favourable druggability score for further comparison of the included metrics: small-molecule drug development (SMB), therapeutic antibody feasibility (FB), safety_(SB), ABability (ABB), new modality (NMB), tissue engagement (TEB). For the Pharos Classification (which reflects the amount of available knowledge for each target) we mapped the different classes (Tclin, Tchem, Tbio, Tdark) to numbers (1.0, 0.75, 0.5, 0.25) to obtain a comparable Pharos classification number (PN).

## Acknowledgements

The results published here are in whole or in part based on data obtained from the AD Knowledge Portal (https://adknowledgeportal.org). The data available in the AD Knowledge Portal would not be possible without the participation of research volunteers and the contribution of data by collaborating researchers. Study data were provided by the Rush Alzheimer’s Disease Center, Rush University Medical Center, Chicago, where data collection was supported through funding by NIA grants P30AG10161, R01AG15819, R01AG17917, R01AG36836, R01AG48015, U01AG46152, the Illinois Department of Public Health (ROSMAP), and the Translational Genomics Research Institute (genomic). The Mayo Clinic Alzheimers Disease Genetic Studies, led by Dr. Nilufer Taner and Dr. Steven G. Younkin, Mayo Clinic, Jacksonville, FL where data collection was supported through funding by NIA grants P50 AG016574, R01 AG032990, U01 AG046139, R01 AG018023, U01 AG006576, U01 AG006786, R01 AG025711, R01 AG017216, R01 AG003949, NINDS grant R01 NS080820, CurePSP Foundation, and support from Mayo Foundation. Study data includes samples collected through the Sun Health Research Institute Brain and Body Donation Program of Sun City, Arizona. The Brain and Body Donation Program is supported by the National Institute of Neurological Disorders and Stroke (U24 NS072026 National Brain and Tissue Resource for Parkinson’s Disease and Related Disorders), the National Institute on Aging (P30 AG19610 Arizona Alzheimer’s Disease Core Center), the Arizona Department of Health Services (contract 211002, Arizona Alzheimers Research Center), the Arizona Biomedical Research Commission (contracts 4001, 0011, 05-901 and 1001 to the Arizona Parkinson’s Disease Consortium) and the Michael J. Fox Foundation for Parkinson’s Research. And, the CommonMind Consortium supported by funding from Takeda Pharmaceuticals Company Limited, F. Hoffmann-La Roche Ltd and NIH grants R01MH085542, R01MH093725, P50MH066392, P50MH080405, R01MH097276, RO1-MH-075916, P50M096891, P50MH084053S1, R37MH057881, AG02219, AG05138, MH06692, R01MH110921, R01MH109677, R01MH109897, U01MH103392, and contract HHSN271201300031C through IRP NIMH. Brain tissue for the study was obtained from the following brain bank collections: the Mount Sinai NIH Brain and Tissue Repository, the University of Pennsylvania Alzheimer’s Disease Core Center, the University of Pittsburgh NeuroBioBank and Brain and Tissue Repositories, and the NIMH Human Brain Collection Core. CMC Leadership: Panos Roussos, Joseph Buxbaum, Andrew Chess, Schahram Akbarian, Vahram Haroutunian (Icahn School of Medicine at Mount Sinai), Bernie Devlin, David Lewis (University of Pittsburgh), Raquel Gur, Chang-Gyu Hahn (University of Pennsylvania), Enrico Domenici (University of Trento), Mette A. Peters, Solveig Sieberts (Sage Bionetworks), Thomas Lehner, Stefano Marenco, Barbara K. Lipska (NIMH)

The results published here are in whole or in part based on data obtained from Agora, a platform initially developed by the NIA-funded AMP-AD consortium that shares evidence in support of AD target discovery. Agora is available at: doi:10.57718/agora-adknowledgeportal.

## Funding Acknowledgement

## Author Contributions

Conceptualization MZ, GS, Data Curation MZ, EML, Formal Analysis MZ, EML, Investigation MZ, EML, Methodology MZ, EML, Resources MZ, EML, Software MZ, Supervision MZ, GS, Validation MZ, Visualization MZ, Writing-original draft MZ, EML, Writing-review & editing MZ, GS.

