## SupplementaryMaterials for "Immediate-Early Genes as Influencers in Genetic Networks and their Role in Alzheimer’s Disease"

### Supplementary Material

#### Supplementary Figures

S1 Mouse

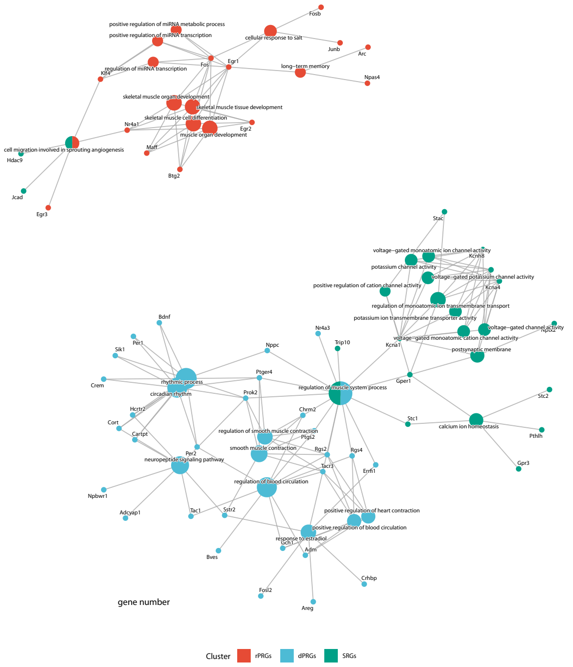


S2 Human

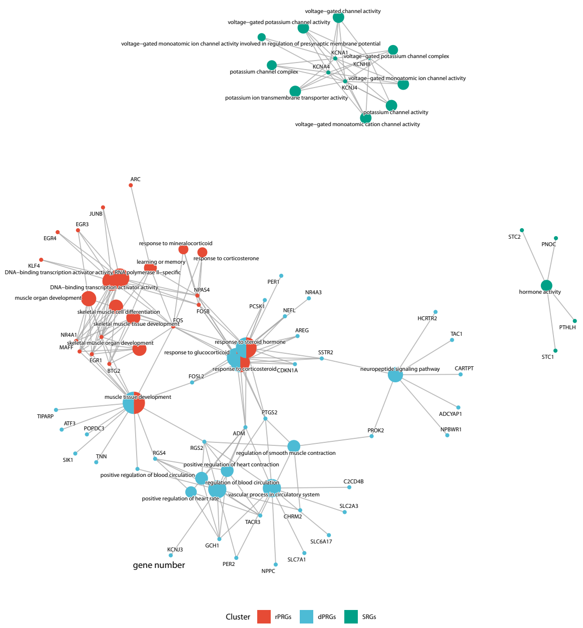


**S1-S2** Bipartite Networks of enriched terms and participating genes per term, colour-coded for each cluster. The size of the circles corresponding to the GO terms indicates the number of genes per term.

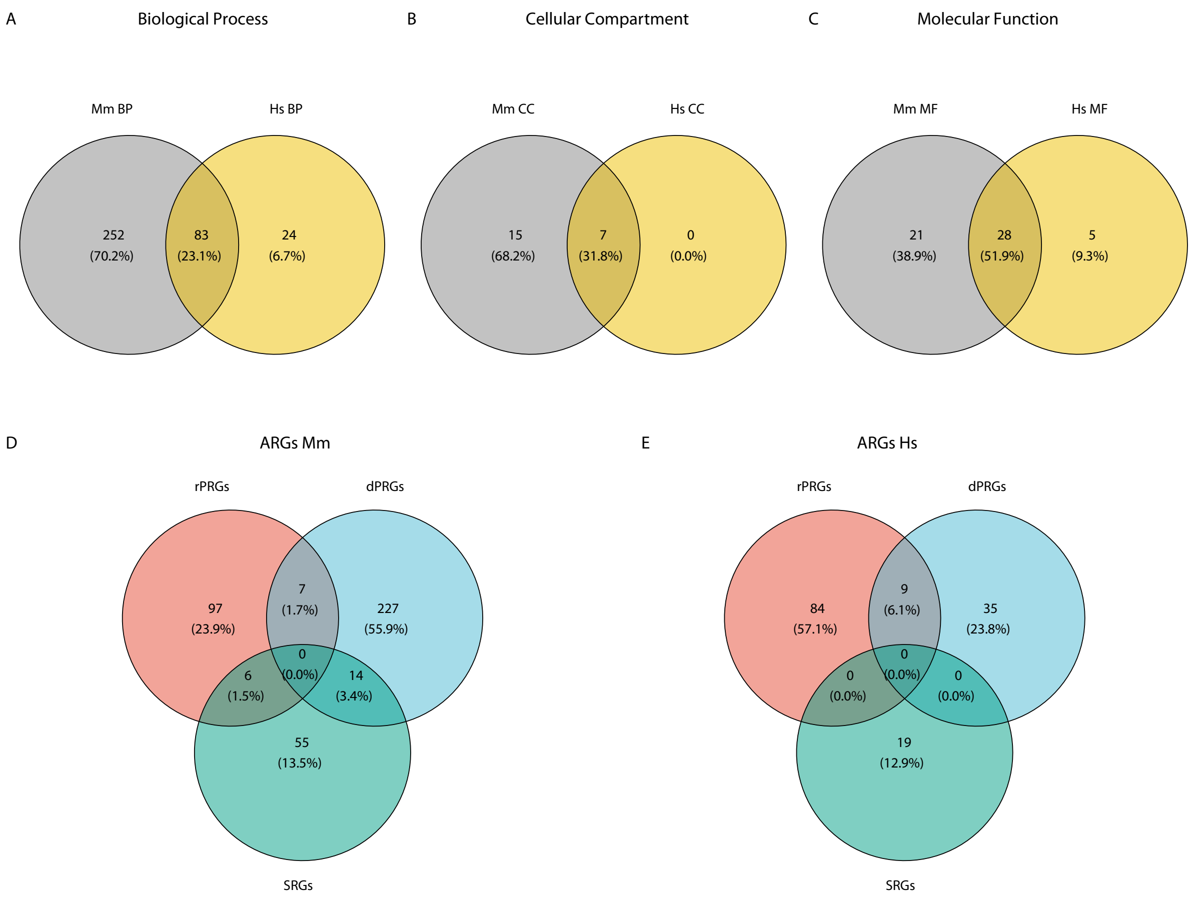


**S3:** Venn diagrams for GO functional analysis.

#### Supplementary Tables

**S1:** Main ARGs table
**S2a:** GO functional analysis Hs
**S2b**: GO functional analysis Mm
**S3:** Random non-ARG pLI and LOEUF
**S4:** Main GWAS results
**S5:** Risk and protective AD variants from the literature
**S6:** ARG-AD borderline protective and ARG-AD borderline risk SNPs
**S7:** LD analysis of *MARK4* variants
**S8:** Degree and IVI centralities for gR-NET-Hs network
**S9:** Degree and IVI centralities for R-NET-Hs network
**S10:** Degree and IVI centralities for AD atlas network
